# Advancing Spatial Pattern Integration in Species Distribution Modelling With Computer Vision: How Grain Size and Extent Concepts Shape Predictions of Emerging Aquatic Insects

**DOI:** 10.64898/2026.10.29.762482

**Authors:** Augustin de la Brosse, Aimi Okabayashi, Thomas Corpetti, Thomas Houet

## Abstract

1. Capturing spatial patterns in ecology necessitates jointly accounting for the concepts of grain size (i.e., the spatial resolution of the input data) and extent (i.e., the area covered by the input data). Species distribution models (SDMs) that rely on traditional machine learning models (e.g., Random Forest, MaxEnt) only integrate the grain size through environmental variables derived from raster data. Recent advances in deep learning have led to the development of Deep SDMs. Unlike traditional SDMs, Deep SDMs directly process image windows representing landscape context. Therefore, they can manipulate both the grain size and extent, through the spatial resolution and size of the windows, respectively.
2. This study evaluates how grain size and window extent affect Deep SDMs, and how grain size alters the importance of variables (i.e., pixels of the image windows). We used multispectral and LiDAR data from drone and satellite acquisitions, with spatial resolutions ranging from 31 cm to 10 m, and varied the window extent between 40 m and 100 m. Grad-CAM was used to assess the importance of variables for each data type and measure their spatial agreement. As a case study, the terrestrial distribution of Plecoptera and Trichoptera, two orders of emerging aquatic insects with contrasting dispersal capacities, was predicted.
3. Results demonstrated the capacity of Deep SDMs to integrate both the grain size and extent. They highlighted substantial variations in performance depending on the grain size (up to 0.26 AUC) and on the extent (up to 0.16 AUC). In particular, the best predictions were obtained using 2.5 m multispectral data at a 40 m extent for Plecoptera (limited dispersers), and the same multispectral resolution combined with LiDAR at a 100 m extent for Trichoptera (greater dispersers). Results also showed a moderate but notable change in variable importance with different grain sizes.
4. Given the growing availability of remote sensing products, our findings advocate for systematic analysis of grain size and extent with Deep SDMs. This is essential to efficiently capture spatio-ecological patterns and to better understand the studied species.

Datasets and code are publicly available^1^.

## 1 Introduction

Species distribution models (SDMs) are essential tools for predicting the spatial distribution of species. Ecologists use them for various purposes, from planning future field surveys for rare species [52] to assessing potential invasive species locations [29]. SDMs predict species distribution by correlating occurrence or abundance data with environmental and spatial variables [11]. Developing such models requires several methodological decisions, including the choice of grain size, defined as the “size of the individual units of observation” [60]. Several studies have analysed how grain size affects SDM performance. Each species responds differently to grain size [9]. However, coarsening the grain size generally tends to degrade model predictive performance [19,39]. An overly coarse grain size could mask certain fine details, despite their importance for some species [53]. An excessively coarse grain size may also overestimate the distribution area [31]. Additionally, the relative importance of environmental variables has also been shown to vary with grain size [9]. Overall, using the wrong grain size can lead to erroneous conclusions. Nevertheless, these studies considered only grain size in the context of traditional SDMs and overlooked the concept of extent (i.e., the area covered by the study), despite also being critical for detecting spatial patterns [60].

Traditional SDMs rely on machine learning (ML) algorithms, such as Random Forest or MaxEnt, which require tabular variables, often derived from remote sensing data. In that setting, the grain size of a variable is implicitly defined by the spatial resolution (the surface covered by a single pixel, determining the level of detail in an image) of the data from which it is extracted. For example, mean NDVI (Normalised Difference Vegetation Index) around a sampling point may be extracted from 10-meter Sentinel-2 or centimetric drone imagery, yielding one tabular value per point with varying spatial detail. Recent advances in deep learning, in particular computer vision techniques such as convolutional neural networks (CNNs), have led to the emergence of Deep SDMs [6,26,55] which have improved predictive performance of SDMs [35]. Unlike traditional SDMs, they use raw remote sensing products. Specifically, they take as input image windows, i.e., square crops of raster data, representing the landscape context around each sampling location [8,58] (Figure 2). This specificity of Deep SDMs has several consequences. Firstly, the grain size is now explicitly defined by the spatial resolution of the image windows. Secondly, it is now possible to integrate and manipulate the concept of extent [60] through the size of the windows, i.e., the ground area covered by the windows. Therefore, since spatial resolution directly affects the performance of deep SDMs, selecting the appropriate remote sensing data is important.

Remote sensing data can be used directly as input or as a derived variable, such as radiometric indices or land cover classifications [2,16,45]. Satellites (e.g., Sentinel-2, Landsat) are regularly used for this purpose [1,2,18,55]. They offer the advantage of high to moderate spatial resolution (10-30 m), rich spectral information, high revisit frequency (several days), and global coverage. Commercial very high-resolution satellites (e.g., Pleiades, SPOT-6) offer great potential when very fine-grained data are needed to model the distribution of small species [12,54]. They offer a finer spatial resolution (up to 30 cm per pixel) than those provided by Sentinel-2 or Landsat. However, their cost and limited availability restrict their use. A low-cost compromise to access very high-resolution data is to use super-resolution (SR) or pan-sharpening algorithms, which improve the spatial resolution of images. Drones also constitute an appealing alternative increasingly used in ecological studies [40,49,3]. Drones have several advantages. They provide very high spatial resolution (up to centimetric) and can be equipped with different types of sensors (multispectral, hyperspectral, thermal, LiDAR, etc). Drones also increase the flexibility of data acquisition (cloud-tolerant, controlled revisit frequency). This is essential to monitor species that undergo seasonal emergence patterns. However, they cover smaller areas than satellites, and their data require more preprocessing than ready-to-use satellite imagery. Overall, there is a wide diversity of remote sensing data types, with trade-offs between grain size, covered extent, and acquisition cost.

To ground this work in a concrete ecological context, this study focuses on emerging aquatic insects. They provide essential ecosystem services to terrestrial ecosystems: additional input of organic matter and nitrogen to surrounding fields [47], contribution to pollination [15], and crop pest control [46]. Therefore, mapping their terrestrial distribution represents an important challenge for mapping these services and understanding aquatic-terrestrial interactions. Among these, we selected Plecoptera and Trichoptera as case studies due to their contrasting spatial distribution behaviours, with Trichoptera dispersing further than Plecoptera [5].

To evaluate the impact of different grain sizes and extents on Deep SDM and assess whether it reflects the contrasting dispersal capacities of Plecoptera and Trichoptera,

1. we analyse the impact of grain size on the predictive performance of convolutional Deep SDMs using operationally relevant remote sensing configurations including drone and satellite data of various spatial resolutions (from 31 cm per pixel to 10 m),
2. we assess the effect of extent by using three different window sizes,
3. we measure the influence of two sensor types, i.e., multispectral and LiDAR,
4. we evaluate whether the grain-size-dependent shift in variable importance, previously observed in traditional SDMs by [9], still holds for Deep SDMs by leveraging the Grad-CAM method [50], an explainable artificial intelligence (XAI) technique adapted to CNNs.

Based on prior research, we hypothesize that the optimal grain size is species-specific and should reflect the life history of the species under study. Consequently, we expect that the distribution of Plecoptera (limited dispersers) will be better predicted by finer-grained data, as the model may capture fine-scale habitat features. Conversely, the distribution of Trichoptera (greater dispersers) will be better predicted by coarser-grained data, as the model may extract landscape-scale patterns. Similarly, we hypothesize that smaller image windows will contain sufficient information to efficiently model the distribution of Plecoptera. In contrast, Trichoptera will necessitate larger windows to yield reliable results.

## 2 Materials and Methods

### 2.1 Materials

#### 2.1.1 Study sites

Experiments were conducted on four agricultural sites in France, each covering approximately 80 ha (Figure 1). They were selected in the European LTSER (Long-Term Socio-Ecological Research) Network along a gradient of agricultural intensity and hedgerow density. The Louroux and Cisse sites (Zone Atelier Loire) represent intensive cropping systems with minimal hedgerows and grasslands. The Timbertière site (Zone Atelier Armorique) combines an intensive livestock/dairy system with a densely wooded landscape. Finally, the Fao site (Zone Atelier Brest-Iroise) features a densely wooded landscape and an extensive dairy farming system.

**Fig. 1.**
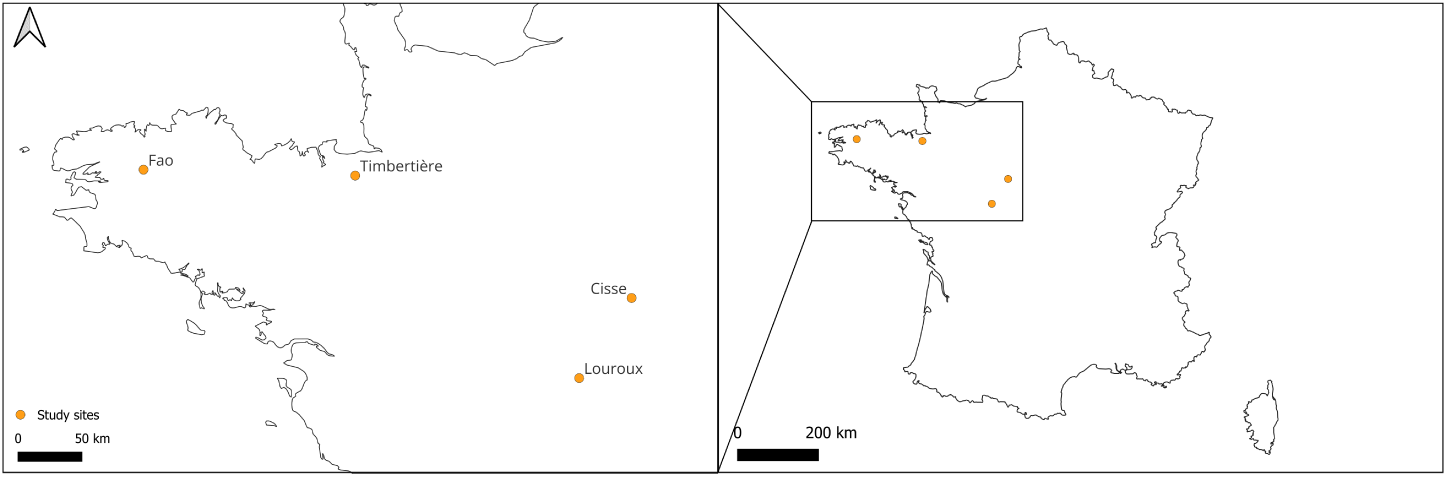
Location of the study sites.

#### 2.1.2 Remotely sensed data

The remote sensing products were selected to represent widely used and emerging data types for Deep SDMs across a range of spatial resolutions, sensor types, and acquisition platforms. The resulting comparisons therefore evaluate operationally relevant data configurations, rather than isolating the effect of grain size and extent under fully controlled conditions.

##### 2.1.2.1 Ultra-fine-grained data (< 1 m/pixel)

**Drone multispectral imagery.** Multispectral (MS) data were acquired using a Trinity F90+ equipped with a MicaSense MX sensor (5 bands). Images for each study site were acquired in April 2024 under homogeneous ceiling conditions. Spectral bands are centred on the blue (475 nm ± 32), green (560 nm ± 27), red (668 nm ± 20), red-edge (717 nm ± 12), and near-infrared (842 nm ± 57) bands. The spatial resolution is 8 cm per pixel for flight at 120 m above ground level (AGL). The orthomosaics were generated for each site / spectral band and radiometrically corrected, accounting for solar irradiance and angle, using a calibrated radiometric target to provide top-of-canopy (TOC) reflectance [4]. Hereafter, this dataset is referred to as “DroneMS31cm”.

**Drone LiDAR imagery.** LiDAR data were acquired on the same day with a Qube240 sensor mounted on Trinity F90+, flying at 120 m AGL, providing a 3D point cloud density of 36.1 points / m^2^ for each site. A digital terrain model (DTM), and a digital surface model (DSM), were derived with a spatial resolution of 10 cm per pixel. Hereafter, this dataset is referred to as “DroneLiDAR31cm”.

Both multispectral and LiDAR data are registered using common ground control points whose positions are set using a GNSS (2-3 cm accuracy). Geometric errors (in X, Y, and Z) are smaller than the pixel size of the data (root mean square error (RMSE) = 7.6 cm XY – 4.1 cm Z). Multispectral and LiDAR-derived data were resampled to 31 cm using linear interpolation in order to limit the computational cost, and are available online [22,23,24,25]

**IGN LiDAR imagery.** DSM and DTM can be downloaded on the French National Institute of Geographic and Forest Information (IGN) platform^2^. The acquisition campaigns were conducted between January and March 2024. DSM and DTM were derived from a point cloud density of at least 10 points/m^2^ and show a 50 cm spatial resolution. Hereafter, this dataset is referred to as “IGNLiDAR50cm”.

##### 2.1.2.2 Very fine-grained data (2.5 m/pixel

**Super-resolution (SR) multispectral imagery.** 2.5-meter SR images were generated from Sentinel-2 data (see Section 2.1.2.3) using a Flow Matching algorithm [33] trained on BreizhSR [42]. The BreizhSR dataset covers the Brittany region in France and pairs Sentinel-2 time series with very high-resolution SPOT-6 imagery (2.5 m, pansharpened) from 2018. The study sites lie within or just beyond the area covered by BreizhSR. This generative approach learns probabilistic transformation pathways between Sentinel-2 and SPOT-6 reference images, while correcting radiometric offsets between sensors to preserve spectral consistency. The output images contain blue, green, red, and near-infrared bands because SPOT-6 provides only these four bands. Implementation details are available in the Appendix. SR remains rarely used to train SDMs, with most applications focused on vegetation mapping [56]. Therefore, this work is an opportunity to better understand the potential benefits of this technique for SDMs. Hereafter, this dataset is referred to as “SRMS2m5”.

**Resampled drone multispectral imagery.** Drone multispectral data are resampled to 2.5 m resolution using linear interpolation. This served two purposes: (1) to create a reference 2.5 m product for comparison with SR imagery, which is inherently imperfect due to algorithmic enhancement, and (2) to approximate the performance of actual high-resolution commercial satellites (e.g., SPOT-6) without requiring their expensive acquisition. Hereafter, this dataset is referred to as “DroneMS2m5”.

##### 2.1.2.3 Fine-grained data (10 m/pixel)

**Sentinel-2 multispectral imagery.** L2A multispectral data were downloaded at 10 m spatial resolution from Copernicus Browser^3^, selecting tiles acquired close to drone survey dates. The acquisition dates for each Zone Atelier (ZA) were the following:

– ZA Brest-Iroise: 18 April 2024;
– ZA Armorique: 20 April 2024;
– ZA Loire: 9 May 2024.

Because drone multispectral data provide only 5 spectral bands, a common subset of Sentinel-2 bands (B02, B03, B04, B05, and B08) is used to isolate grain size and extent effects from spectral composition effects. Hereafter, this dataset is referred to as “SatelliteMS10m”.

#### 2.1.3 Species distribution data

Information on presence and absence was collected using 196 sticky traps deployed along the 4 study sites for two weeks in April 2024, during the emergence period. Specifically, 50 sticky traps were installed in the Louroux, Cisse, and Timbertière sites and 46 in the Fao site, all following the same protocol described in [17].

### 2.2 Methods

This section outlines the methodology employed to analyse the effect of grain size and extent for Deep SDMs. First, the different experiments are described. Then, the model architectures and training setup are presented. Next, the algorithm used to analyse the importance of variables is detailed. Note that when the input is an image, the variables are the pixels of the input image. Finally, procedures used to evaluate the predictive performance of the Deep SDMs and changes in variable importance are explained.

#### 2.2.1 Experimental protocol

##### 2.2.1.1 Tested configurations

Fourteen different data combinations were tested to predict the spatial distribution of Plecoptera and Trichoptera: 6 single-sensor configurations and 8 dual-sensor configurations, each combining one multispectral data type (drone, resampled drone, super-resolution, or Sentinel-2) with one LiDAR data type (drone or IGN). Table 1 summarizes these combinations. These configurations enable the evaluation of the respective impact of grain size and sensor type on predictions.

**Table 1.** Summary of the data combinations tested.

| Sensor type | Multispectral |  |  |  | LIDAR |  |
| --- | --- | --- | --- | --- | --- | --- |
|  | DroneMS<br>31cm | DroneMS<br>2m5 | SRMS<br>2m5 | Satellite<br>MS10m | Drone<br>LIDAR31cm | IGN<br>LIDAR50cm |
| Config 1 | X |  |  |  |  |  |
| Config 2 |  | X |  |  |  |  |
| Config 3 |  |  | X |  |  |  |
| Config 4 |  |  |  | X |  |  |
| Config 5 |  |  |  |  | X |  |
| Config 6 |  |  |  |  |  | X |
| Config 7 | X |  |  |  |  | X |
| Config 8 | X |  |  |  | X |  |
| Config 9 |  | X |  |  |  | X |
| Config 10 |  | X |  |  | X |  |
| Config 11 |  |  | X |  |  | X |
| Config 12 |  |  | X |  | X |  |
| Config 13 |  |  |  | X |  | X |
| Config 14 |  |  |  | X | X |  |

##### 2.2.1.2 Creation of the 40 × 40 m, 70 × 70 m and 100 × 100m image windows

For each configuration, three different window extents (40, 70 and 100 m) were tested by extracting from the full raster 3 windows per sampling point and data type (Figure 2). Therefore, the number of pixels per window varied by data source: for example, at 40 m, DroneMS31cm and SatelliteMS10m windows contain 128 × 128 and 4 × 4 pixels, respectively. This allows us to assess the impact of extent on Deep SDMs.

**Fig. 2.**
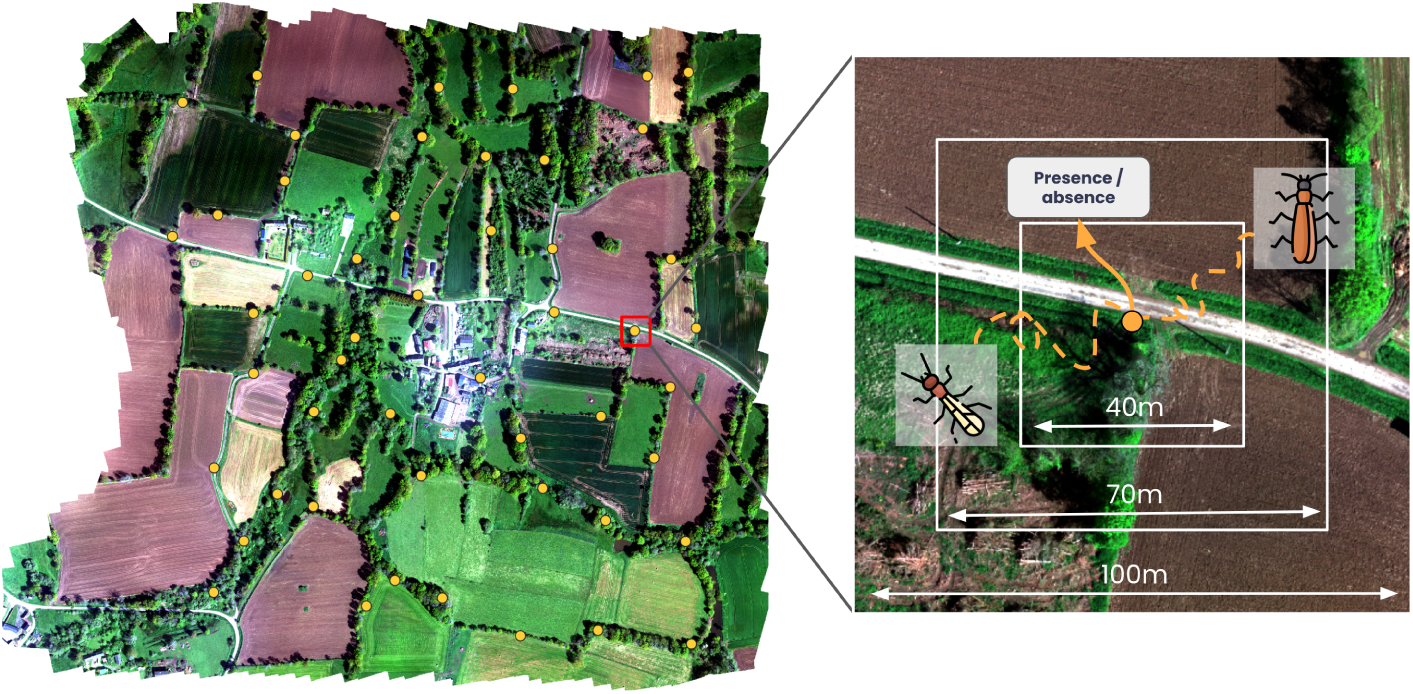
*Left panel*: global overview of the Timbertière acquisition site. *Right panel*: Illustration of the 40 *×* 40 m, 70 *×* 70 m and 100 *×* 100 m windows which were used as input to the model. These windows are centred on the sticky trap, which gives information on the presence and absence of Plecoptera and Trichoptera.

#### 2.2.2 Deep SDMs

##### 2.2.2.1 Architecture

The Deep SDMs used in this study are based on CNNs. Such models are divided into 2 parts: a feature extractor that extracts the most meaningful information from the input data to best perform the desired task, and a classification head that outputs the final prediction.

We use two different feature extractors based on the spatial resolution of the data. When data are ultra-fine-grained (see Section 2.1.2), we use the multi-branch extractor, CerberusCNN, described in [7], which obtains reliable results for both Plecoptera and Trichoptera while being lightweight regarding the number of parameters. However, this architecture is not adapted to coarser-resolution data (very-fine-grained and fine-grained data) as the kernel size (up to 7×7 pixels) may exceed the input window dimensions. Kernels are fundamental components of CNNs: they are small matrices of learnable weights that are convolved across the input to detect local patterns (e.g., edges, textures, or spectral signatures). Their spatial dimensions must be smaller than the input to enable the extraction of local features. Therefore, we used a similar multi-branch extractor, TinyCerberusCNN, adapted to low-resolution data. The three branches (1×1 and 3×3 convolutions, plus global pooling) capture pixel-level, local, and window-level information. Both feature extractors produce a 192-dimensional vector. Note that using the low-resolution feature extractor for high-resolution data degraded performance (see Appendix for more details). The classification head is a multilayer perceptron with a dense layer, a dropout layer, and a final dense layer with sigmoid activation for binary presence/absence prediction.

Depending on the tested configuration, its input is either a 192-dimensional vector (single-sensor configurations) or a 384-dimensional vector (dual-sensor configurations). An exhaustive description of the architectures is available in the Appendix.

##### 2.2.2.2 Training setup

Data leakage, a common issue in SDMs [39], occurs when a model leverages information from the test dataset during training. This creates an illusion of high performance during training, but the model fails to generalise to real-world predictions. As spatially close windows may overlap and cause data leakage, rasters were split independently for each site based on longitude before extracting the windows. For each site, the easternmost 20% (41 occurrences) were used for testing, the next 20% (31 occurrences) were used for validation, and the remaining samples (124 occurrences) were used for training. The label distribution is displayed in Table 2. Regarding preprocessing, the no-data values of the windows were replaced with 0, and then min-max normalised per band. To increase the size of our training dataset and improve the regularisation of the models, we applied data augmentation techniques. This included weak random transformations (horizontal and vertical flips, rotations) and MixUp [61], a method that creates new synthetic samples by linearly combining pairs of images and their corresponding labels.

**Table 2.** Presence rate for each taxon across the training, validation, and test splits. Example: the presence rate of Plecoptera in the train set is 45.2%.

| Taxon | Train (n = 124) | Validation (n = 31) | Test (n = 41) |
| --- | --- | --- | --- |
| Plecoptera | 45.2% | 38.7% | 36.6% |
| Trichoptera | 37.1% | 38.7% | 46.3% |

All models were trained using a binary cross-entropy loss [38] and an AdamW optimizer with a learning rate of 0.001 and a weight decay of 0.0001 [36]. We set the batch size to 8 and trained models for up to 50 epochs, with early stopping triggered after 5 epochs without improvement of the loss on the validation set.

#### 2.2.3 Analysing variable importance with XAI

As noted by [9], the importance of variables changes according to grain size in traditional SDMs. The variable permutation method used by [9] is not suitable for Deep SDMs that take images as input. Therefore, we used Grad-CAM (Gradient-weighted Class Activation Mapping) [50], a widely used XAI method for images. It generates an activation heatmap (where activation refers to the strength of the response of a model to specific input regions) that highlights the variables (i.e., the pixels of the image window) that most influenced the model prediction. To achieve this, it weights the feature maps (i.e., the output) of the final convolutional layer with the gradients of the target class and sums the results to produce a coarse localization map. As the architectures are multi-branch, we generate a heatmap for the final convolutional layer of each branch and average them (after normalising and resizing them to a common size). The heatmaps are produced separately for each data type at a common spatial extent of 100 m to ensure that the explanation remains specific to each data type (Figure 4). Then, spatial agreement was measured between the different heatmaps.

#### 2.2.4 Evaluation metrics

##### 2.2.4.1 Distribution predictions

The models were evaluated based on the Area Under the Curve (AUC) metric, which is frequently used for SDMs [17]. A model is deemed reliable when its AUC exceeds 0.70 and excellent when it surpasses 0.80 [20]. We trained each model three times to account for the inherent variability of deep learning models, similarly to [7].

##### 2.2.4.2 Comparison of variable importance based on the grain size

The heatmaps representing the importance of variables were compared both qualitatively (visual inspection) and quantitatively. For the quantitative analysis, we evaluated changes in variable importance using Pearson’s correlation coefficient between pairs of data types for samples for which both models correctly predicted presence (true positive) in the test dataset. This ensured that both heatmaps explained the same ecological outcome. However, as grain size varies widely across data types, we divided each heatmap into a 10×10 cell grid to align with the coarsest grain size (SatelliteMS10m). We then averaged activation values within each cell and calculated the Pearson correlation between the binned activations of each pair of data types across all shared true-positive samples. Higher correlation indicates stronger agreement in the spatial regions considered important by both models. This approach ensures a fair comparison of spatial focus between data types, regardless of their original resolution.

## 3 Results

### 3.1 Distribution predictions

Figure 3 shows important differences in performance depending on the grain size, the extent, and the data configuration used.

**Fig. 3.**
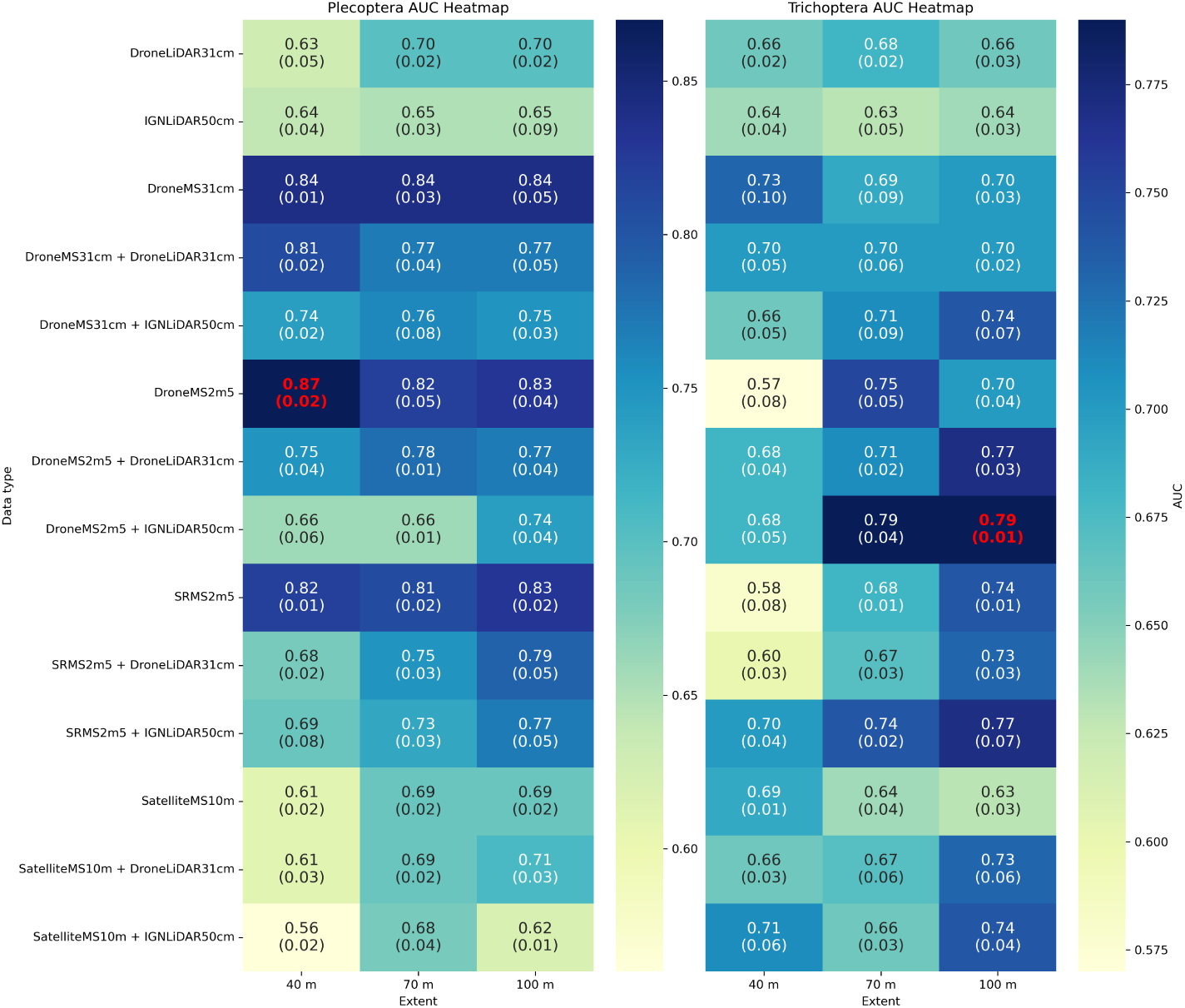
Model predictive performance per taxon: mean AUC and standard deviation over three training runs. Best scores are in red. The x-axis represents window extents, and the y-axis represents data types.

For Plecoptera, single-sensor models using ultra-fine and very fine-grained multispectral data achieve high performance (at least 0.82 AUC) regardless of the window extent. However, performance drops to a maximum of 0.69 AUC with Sentinel-2 data. Conversely, single-sensor models using LiDAR fail to predict the distribution of both taxa reliably, with AUC below 0.7 (except for drone LiDAR with a window extent of 100 m). Furthermore, adding LiDAR to multispectral data is detrimental to the model’s performance. On average over all window extents, it degrades the performance by 0.07 points.

For Trichoptera, although LiDAR cannot reliably predict its distribution, adding it to multispectral data improves performance by 0.03 points on average across all spatial extents. Specifically, IGNLiDAR50cm (acquired in the winter preceding the emergence) increases AUC more than DroneLiDAR31cm (acquired during the emergence in spring). Also, the highest performances are observed at the 100 m window extent.

Overall, the best performance is obtained using the DroneMS2m5 alone with a window extent of 40 m for Plecoptera (0.87 AUC). For Trichoptera, it is obtained when combined with IGNLiDAR50cm data with a window extent of 100 m (0.79 AUC).

### 3.2 Comparison of variable importance based on the grain size

Figure 4 shows an example of variable importance by data type and taxon. We see in these examples that the highlighted regions are very different when using LiDAR and multispectral data. LiDAR-based models tend to show very specific regions, while multispectral-based SDMs show large areas, mostly forest. As expected, the coarser the grain size, the coarser the heatmap.

**Fig. 4.**
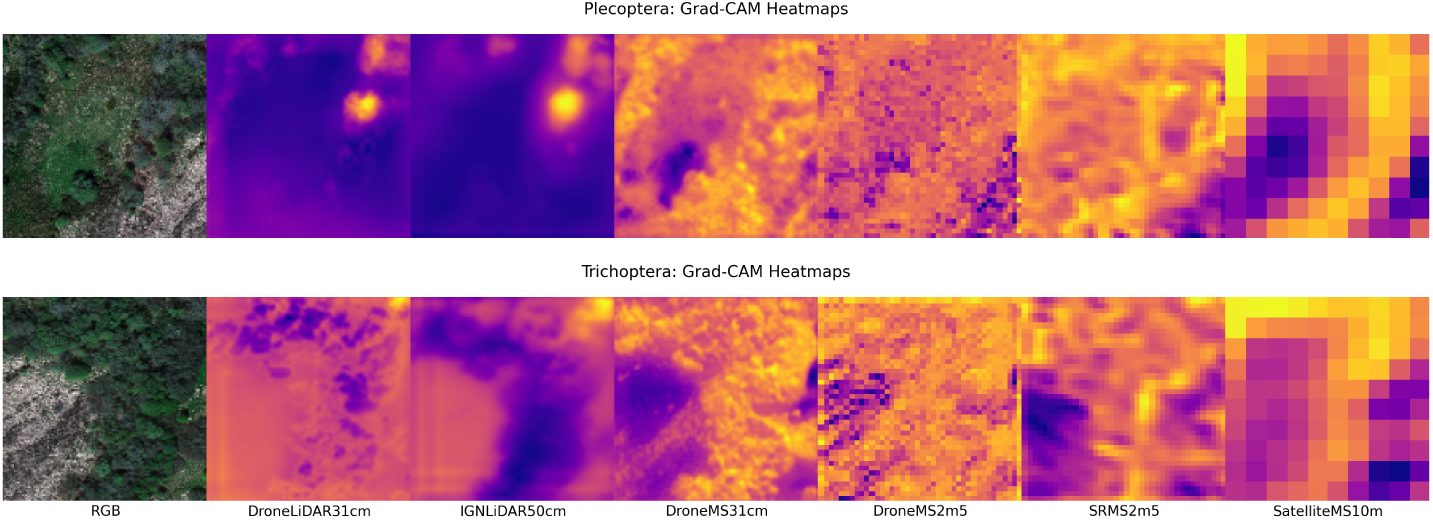
Examples of Grad-CAM heatmaps for Plecoptera and Trichoptera. All models predicted presence. The colour yellow signifies a highly influential region for the prediction of presence (and vice versa with purple).

Figure 5 reveals the global spatial agreement between Grad-CAM heatmaps across data types when predicting species presence. For both Plecoptera and Trichoptera, the LiDAR heatmaps show low or no correlation with the multispectral heatmaps, with Pearson’s coefficients typically ranging from −0.2 to 0.3. Similarly, low correlations (r < 0.4) between drone LiDAR and IGN LiDAR indicate low spatial agreement even within LiDAR data. In contrast, multispectral heatmaps demonstrate a moderately positive correlation, with coefficients mostly between 0.4 and 0.7.

**Fig. 5.**
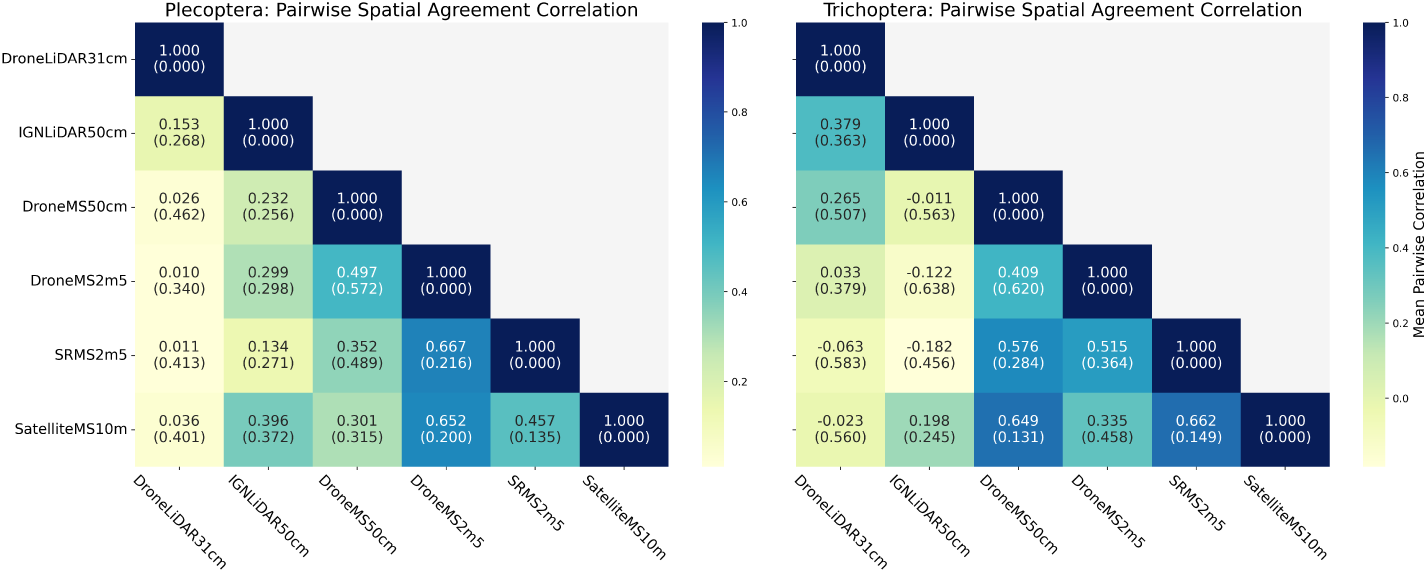
Pairwise spatial agreement correlation as measured by Pearson’s coefficient for Plecoptera and Trichoptera.

## 4 Discussion

### 4.1 Grain size and extent must be integrated into the experimental design of Deep SDM development

**The concepts of grain size and extent are inherently linked in ecology and must be considered jointly to enable a holistic exploration of scale effects** [43,60]. Indeed, grain size controls the details of landscape patterns throughout pixels, whereas window extent integrates the influence of the surrounding landscape for each sample. However, the environmental variables that traditional SDMs use as input do not account for the extent. Consequently, the effect of the grain size on SDMs could not be assessed alongside the effect of extent. By processing image windows as input, computer vision algorithms such as CNNs enable these two dimensions to be manipulated to fully account for scaling effects. This capacity allows Deep SDMs to effectively capture spatial patterns. Our results confirm their critical role: grain size and window extent must be treated as key experimental factors to capture spatial patterns of landscape. **Focusing on only one dimension may lead to misleading conclusions about species-environment relationships and model robustness.**

**The characteristics (grain size, extent, sensor type) of Deep SDM input data must reflect species ecology**, as illustrated by the contrasting responses of Plecoptera and Trichoptera. The former have moderate dispersal capacities. 50% of Plecoptera do not disperse further than 3 m and only 10% can be found over 100 m from their emergence sources [5]. This spatially restricted distribution aligns with the greater performance of the smallest window extent (40 m). Any landscape information beyond this is probably less informative. Also, the best results are obtained with an intermediate grain size (DroneMS2m5), indicating that mesoscale details may be optimal for explaining their distribution. One possible explanation is that finer-grained data introduce unnecessary variations (e.g., shadows), slightly degrading model quality, as seen in previous studies [34,41].

Conversely, coarser-grained data are insufficient for the model to capture relevant details and to provide reliable predictions. Finally, LiDAR data often reduce performance compared to multispectral-only models, meaning that all the information needed to describe the dispersion of Plecoptera is included in the multispectral data. These outcomes corroborate those found by [17]. They observed (1) that variables aggregated over 25 m have no positive influence on the improvement of their models, and (2) that elevation variables do not stand out as main driving factors.

Trichoptera have greater dispersal capacities (potentially over 300 m from their emergence source in agricultural landscapes) [5]. This may explain why using a wider window extent (100 m) leads to better model performance, with smaller extents providing incomplete landscape information related to their dispersal. Moreover, similarly to Plecoptera, mesoscaled information (DroneMS2m5) appears to be optimal for predicting the distribution of Trichoptera. The positive contribution of LiDAR to the quality of models suggests that topography (ground and surface) contains complementary information for explaining the distribution of Trichoptera. This is consistent with the study by [27], which indicated that the abundance of several Trichoptera species was significantly correlated with the surface height. Notably, IGNLiDAR50cm, acquired before the emergence period (January-March 2024), provided better results than DroneLiDAR31cm, acquired during the emergence period (April-May 2024). This aligns with the results found by [14], who demonstrated that Trichoptera emergence in spring is strongly determined by wet-season (October-March) hydrologic conditions (e.g., rainfall, peak discharge). These conditions are directly tied to winter topography, which LiDAR describes. The findings outlined above confirm the work of [32], namely that there is no single scale, including grain and extent, at which ecological phenomena can be studied. Each organism perceives its environment differently, depending on its life-cycle strategy (e.g., its dispersal). It is therefore necessary to calibrate the scale parameters of the study to the organism in question in order to fully understand its behaviour.

Contrary to previous studies related to traditional SDMs, which found existing but limited effects [19,39,51], **we see massive changes when using data with different grain sizes to train Deep SDMs** (e.g., a 0.26 AUC difference between the Plecoptera model using SatelliteMS10m at 40 m and the model using DroneMS2m5 also at 40 m). However, similarly to traditional SDMs [9], our study demonstrates that the finest grain size does not always provide the best distribution predictions, suggesting an optimal grain size for each taxon. Super-resolution data performed reliably, supporting their potential for Deep SDMs, as already noted by [56]. Indeed, it is more cost-effective than commercial high-resolution satellites and is more suited to large areas than drones. **The window extent is also responsible for important performance variations**. For instance, Trichoptera models show a 0.16 AUC difference between 40 m and 100 m window extents despite both using SRMS2m5. The window is an important factor enabling Deep SDMs to capture the influence of the landscape structure on species distribution around the sample [10]. Its extent should therefore be adapted to align with the species of interest. Finally, our study showed that adding more data is not always relevant. Indeed, the combination of LiDAR with multispectral data resulted in models that were less effective at predicting the distribution of Plecoptera. Instead, one must focus on identifying the appropriate data types. This confirms the findings of [37]. They showed that coupling variables selected based on expert knowledge with ready-to-use climate variables was detrimental to the performance of traditional SDMs compared to using only well-defined expert variables.

A possible next step is to leverage the flexibility of deep learning models to train multi-grain-size models, similar to multi-scale models [57] that integrate image windows of varying extents as input. This could enable Deep SDMs to automatically identify the optimal grain size for each species, thereby improving model robustness across heterogeneous environments.

### 4.2 Moderate but notable variable importance changes across grain sizes

Our Grad-CAM heatmap analysis verifies for Deep SDMs what [9] found for traditional SDMs: **the importance of variables, the pixels of the input image window, changes as grain size varies**. This means that depending on the grain size, Deep SDMs will not focus on the same regions in the input image window. However, this spatial focus is not completely different. Deep SDMs using multispectral data examine approximately the same regions, as indicated by moderate spatial agreement among the heatmaps generated by these models. This is likely because multispectral data capture similar spectral signatures of vegetation and land cover, regardless of the grain size. In contrast, there is no correlation between the Grad-CAM heatmaps generated by the Deep SDMs using the two LiDAR data types, meaning that these models look at very distinct regions. A possible explanation is the different acquisition periods of IGN LiDAR (winter) and drone LiDAR (spring), unlike multispectral data. This results in different representations of the landscape. This approach enabled us to verify how variable importance changes with grain size variation. However, it seems challenging to extract relevant global ecological knowledge about Plecoptera and Trichoptera without fine-grained land cover maps for heatmap comparison. Here, we could only understand each prediction individually. A possible alternative approach is the use of concept-based XAI [7]. This approach aims to explain AI models by quantifying the influence of high-level, interpretable concepts (e.g., wheat crop, forest) on the predictions.

### 4.3 Rethinking the development and use of Deep SDMs

Overall, our study on Plecoptera and Trichoptera demonstrates that analysing the effects of grain size and extent yields more than simply improved predictions. In fact, **systematically varying grain size and window extent serves as a powerful diagnostic tool for improving the understanding of ecological processes and dynamics**. First, as the optimal configuration is rarely obvious a priori, even with expert knowledge [44,60], this approach provides a robust data-driven way to identify it for each species. Second, conducting an extensive analysis of these parameters yields insights that go beyond maximising predictive accuracy. It deepens our understanding of the species under study. For example, by varying the window extent, we identified the most meaningful extent to study the distribution of Plecoptera and Trichoptera [32]. This framework also generates new ecological hypotheses, such as the influence of environmental conditions in the months preceding emergence on the spatial distribution of emerging aquatic insects.

### 4.4 Limitations and perspectives

This study focuses on remote sensing products that are operationally relevant for Deep SDMs. For example, Sentinel-2 data are extensively used to model the distribution of both fauna [13,45] and flora [30]. Similarly, drone-acquired data is also widely used as input for SDMs [21,28,59]. These products span various spatial resolutions commonly encountered in practical SDM workflows, which makes our study particularly valuable for SDM developers. However, the different data come from different remote sensing platforms. Therefore, the effect of grain size is not fully isolated, although the bands were chosen in order to reduce sensor effects and the acquisition dates were selected to minimise the temporal effect. A perspective on this study is using solely drone data and resampling it to different spatial resolutions. That way, temporal or spectral effects would be limited to isolate the grain size effect.

Another perspective is the integration of the temporal resolution with the other parameters. Indeed, satellites and drones can provide time series, opening new perspectives for SDMs. Depending on their temporal resolution (frequency) and depth (length of time-series), they could contribute to improving the ability of SDMs to capture spatio-temporal ecological processes.

## Acknowledgements

This study was funded by the French National Research Agency, STRANGE project (ANR-23-CE03-0006). It has also benefited from the logistical support of the Zone Atelier Loire, Zone Atelier Armorique, and Zone Atelier Brest-Iroise. We would like to also thank Thibaut Peres, Benjamin Bergerot and Laura Pellan for their precious contribution to the collection of field data.

## Author contributions

A. de la Brosse, T. Corpetti and T. Houet conceived the ideas and designed the methodology; T. Houet, A. de la Brosse and A. Okabayashi collected the data; A. de la Brosse analysed the data; A. de la Brosse led the writing of the manuscript. All authors contributed critically to the drafts and gave final approval for publication. Our study brings together authors from different academic backgrounds, including ecology, geography, remote sensing and data science. All authors were engaged early on with the research and study design to ensure that the diverse set of perspectives they represent was considered from the onset. The team of authors is not gender balanced.

## Authors’ Declaration

All authors on the manuscript have seen and approved the submitted version of the manuscript, have substantially contributed to the work, and all persons entitled to co-authorship have been included.

## Data availability statement

– Datasets are available on a Zenodo repository: https://zenodo.org/records/23234877.
– Code is available on a GitHub repository: https://anonymous.4open.science/r/grain-size-extent-deep-sdm-8251/.

## AI Usage Declaration

The authors used the GitHub Copilot AI assistant to improve the readability and grammar of the manuscript and enhance the clarity of the code (docstrings).

## A More details about the feature extractors

### A.1 Exhaustive description of the feature extractors

#### A.1.1 CerberusCNN feature extractor

This architecture, described in [7], has a three-branch convolutional design to extract multi-scale spatial features. The three branches process the input separately through two convolutional blocks with kernel sizes of 3×3, 5×5 and 7×7, respectively. The first block includes a max-pooling layer with sizes of 2×2, 4×4 and 8×8, while the second block applies a max-pooling layer of size 2 across all branches. A batch normalisation layer separates these two blocks. The extracted features are then aggregated using 2D global average pooling, concatenated, and passed through the classification head. This feature extractor contains approximately 180k trainable parameters.

#### A.1.2 TinyCerberusCNN feature extractor

The low-resolution extractor adopts the same three-branch philosophy as CerberusCNN but replaces large-kernel convolutions with operations suited to the coarser input, since kernels of 5 or 7 pixels would exceed the patch dimensions at this resolution. The first branch captures pixel-level spectral signatures using a 1×1 convolution, the second branch captures local neighborhood patterns using a 3×3 convolution, and the third branch captures the global patch context by directly applying global average pooling to the input, followed by a dense layer. Each branch is followed by a batch normalisation layer and reduced via 2D global average pooling (except the third branch, which is pooled before the dense layer). The extracted features are then concatenated and passed through the classification head, in the same way as CerberusCNN. This design produces an output of 192 features, matching the dimensionality of the high-resolution extractor. This feature extractor contains approximately 4k trainable parameters.

### A.2 Suitability of TinyCerberusCNN for ultra-fine-grained data

We compared the performance of TinyCerberusCNN, designed for coarser data, against CerberusCNN on the ultra-fine-grained data (DroneMS31cm, DroneLi-DAR31cm, IGNLiDAR50cm) to justify the use of two distinct feature extractors depending on grain size (see Section 2.2.2.1). We restricted this comparison to the 40 m window extent, as results at this extent were already conclusive.

#### A.2.1 Results

Results displayed in Table 3 demonstrate that the CerberusCNN feature extractor consistently provides better results than TinyCerberusCNN. For Plecoptera, there is a 0.07 average AUC difference between the CerberusCNN and the TinyCerberusCNN, while for Trichoptera, the difference is 0.04. The gap was largest for the two LiDAR products and comparatively small for multispectral data. It suggests that TinyCerberusCNN’s smaller receptive field is particularly limiting when extracting structural information from LiDAR-derived elevation data.

**Table 3.** Model predictive performance per taxon and feature extractor: mean AUC and standard deviation over three training runs (40 m window extent). Best scores are in bold. The difference is calculated as CerberusCNN - TinyCerberusCNN.

| Taxon | Plecoptera |  |  | Trichoptera |  |  |
| --- | --- | --- | --- | --- | --- | --- |
|  | TinyCerberus<br>CNN | CerberusCNN | Diff. | TinyCerberus<br>CNN | CerberusCNN | Diff. |
| DroneMS31cm | 0.82 (0.01) | **0.84 (0.01)** | 0.02 | 0.68 (0.07) | **0.73 (0.10)** | 0.05 |
| DroneLiDAR31cm | 0.56 (0.03) | **0.63 (0.05)** | 0.07 | 0.61 (0.04) | **0.66 (0.02)** | 0.05 |
| IGNLiDAR50cm | 0.53 (0.06) | **0.64 (0.04)** | 0.11 | 0.63 (0.06) | **0.64 (0.04)** | 0.01 |

#### A.2.2 Discussion

These results confirm that TinyCerberusCNN, whose smaller kernels (1×1 and 3×3) are designed to better exploit the limited pixel dimensions of coarse-grained patches, is not well suited to ultra-fine-grained data. Conversely, the larger receptive fields (up to 7×7) of CerberusCNN can better exploit the higher dimensionality of the data induced by the additional spatial details. Using TinyCerberusCNN on this data would therefore underestimate the true predictive potential of ultra-fine-grained products and risk biasing our grain-size comparisons. This confirms our choice to pair each grain size with its most suitable feature extractor, consistent with standard practice in real-world Deep SDM applications, where model architecture is itself an object of optimisation.

## B Implementation details of the Flow Matching algorithm

The architecture that was used to generate the Super-Resolution images is a U-Net [48] with 50M parameters. The Flow Matching implementation details are available on this GitHub repository: https://github.com/facebookresearch/flow_matching. Gaussian Noise (std = 2) was added to the Sentinel-2 images.

## Footnotes

1 Please refer to Data Availability section.

2 https://cartes.gouv.fr/telechargement/IGNF_NUAGES-DE-POINTS-LIDAR-HD

3 https://browser.dataspace.copernicus.eu

## References

1. Ahmadi, K., Mahmoodi, S., Pal, S.C., Saha, A., Chowdhuri, I., Nguyen, T.T., Jarvie, S., Szostak, M., Socha, J., Thai, V.N.: Improving species distribution models for dominant trees in climate data-poor forests using high-resolution remote sensing. Ecological Modelling 475, 110190 (2023). 10.1016/j.ecolmodel.2022.110190

2. Ahmed, N., Atzberger, C., Zewdie, W.: Species distribution modelling performance and its implication for sentinel-2-based prediction of invasive prosopis juliflora in lower awash river basin, ethiopia. Ecological Processes 10(1), 18 (2021)

3. Alvarez-Vanhard, E., Houet, T., Mony, C., Lecoq, L., Corpetti, T.: Can uavs fill the gap between in situ surveys and satellites for habitat mapping? Remote Sensing of Environment 243, 111780 (2020)

4. Assmann, J.J., Kerby, J.T., Cunliffe, A.M., Myers-Smith, I.H.: Vegetation monitoring using multispectral sensors—best practices and lessons learned from high latitudes. Journal of Unmanned Vehicle Systems 7(1), 54–75 (2018)

5. Bergerot, B., Piscart, C., Roussel, J.: Tightly intertwined: Waterscapes prompt urgent reconsideration of aquatic insects and their role in agricultural landscapes. Science of the Total Environment 966, 178728 (2025)

6. Botella, C., Joly, A., Bonnet, P., Monestiez, P., Munoz, F.: A deep learning approach to species distribution modelling. In: Multimedia tools and applications for environmental & biodiversity informatics, pp. 169–199. Springer (2018)

7. de la Brosse, A., Garreau, D., Houet, T., Corpetti, T.: A high-resolution landscape dataset for concept-based xai with application to species distribution models. In: Joint European Conference on Machine Learning and Knowledge Discovery in Databases. pp. 619–635. Springer (2026)

8. Civco, D.L., Hurd, J.D., Wilson, E.H., Arnold, C.L., Prisloe, MP, J.: Quantifying and describing urbanizing landscapes in the northeast united states. Photogrammetric Engineering and Remote Sensing (2002)

9. Connor, T., Hull, V., Viña, A., Shortridge, A., Tang, Y., Zhang, J., Wang, F., Liu, J.: Effects of grain size and niche breadth on species distribution modeling. Ecography 41(8), 1270–1282 (2018)

10. Deneu, B., Servajean, M., Bonnet, P., Botella, C., Munoz, F., Joly, A.: Convolutional neural networks improve species distribution modelling by capturing the spatial structure of the environment. PLoS computational biology 17(4), e1008856 (2021)

11. Elith, J., Leathwick, J.R.: Species distribution models: ecological explanation and prediction across space and time. Annual review of ecology, evolution, and systematics 40(1), 677–697 (2009)

12. Espel, D., Courty, S., Auda, Y., Sheeren, D., Elger, A.: Submerged macrophyte assessment in rivers: An automatic mapping method using pléiades imagery. Water Research 186, 116353 (2020)

13. Estopinan, J., Servajean, M., Bonnet, P., Munoz, F., Joly, A.: Deep species distribution modeling from sentinel-2 image time-series: a global scale analysis on the orchid family. Frontiers in Plant Science 13, 839327 (2022)

14. Feminella, J.W., Resh, V.H.: Hydrologic influences, disturbance, and intraspecific competition in a stream caddisfly population. Ecology 71(6), 2083–2094 (1990)

15. Figueroa, J.M.T.d., Sánchez-Ortega, A.: Imaginal feeding of twelve nemouroidean stonefly species (insecta, plecoptera). Annals of the Entomological Society of America 93, 251–53 (2000)

16. Friedl, M.A., Sulla-Menashe, D., Tan, B., Schneider, A., Ramankutty, N., Sibley, A., Huang, X.: Modis collection 5 global land cover: Algorithm refinements and characterization of new datasets. Remote sensing of Environment 114(1), 168–182 (2010)

17. Gerber, R., Piscart, C., Roussel, J.M., Georges, R., Houet, T., Royer, J., Bergerot, B.: Landscape models can predict the distribution of aquatic insects across agricultural areas. Landscape Ecology 38(11), 2917–2929 (2023)

18. Giachello, S., Lefosse, S., Simoncini, A., Bonato, L.: Species distribution models for the conservation of a micro-endemic animal: the contribution of regional land cover. Biodiversity and Conservation 34(5), 1689–1707 (2025)

19. Guisan, A., Graham, C.H., Elith, J., Huettmann, F., Group, N.S.D.M.: Sensitivity of predictive species distribution models to change in grain size. Diversity and distributions 13(3), 332–340 (2007)

20. Guisan, A., Thuiller, W., Zimmermann, N.E., Di Cola, V., Georges, D., Psomas, A.: Habitat suitability and distribution models. Habitat suitability and distribution models (2017)

21. Habel, J.C., Teucher, M., Ulrich, W., Bauer, M., Rödder, D.: Drones for butterfly conservation: larval habitat assessment with an unmanned aerial vehicle. Landscape Ecology 31(10), 2385–2395 (2016)

22. Houet, T., Peres, T., de la Brosse, A.: La cisse 2024-04-24 - dataset drone (2026). 10.35110/1dd6c296-c6cd-4514-b581-88d9cad64b7b

23. Houet, T., Peres, T., de la Brosse, A.: La timbertiere - dataset drone - 2024-04-24&29 (2026). 10.35110/fc45bf4f-7479-47d5-a05a-95d868caf14b

24. Houet, T., Peres, T., de la Brosse, A.: Le fao - dataset drone - 2024-04-18 (2026). 10.35110/79402bee-14cf-4e2c-84c3-8238a44edbd2

25. Houet, T., Peres, T., de la Brosse, A.: Le louroux 2024-04-24 - dataset drone (2026). 10.35110/75a44ad8-ab96-4bc3-aaa9-4ff7b5ac65c9

26. Hu, Y., Si-Moussi, S., Thuiller, W.: Introduction to deep learning methods for multi-species predictions. Methods in Ecology and Evolution 16(1), 228–246 (2025)

27. Jackson, J.K., Resh, V.H.: Distribution and abundance of adult aquatic insects in the forest adjacent to a northern california stream. Environmental entomology 18(2), 278–283 (1989)

28. Kim, Y., Lee, S., Shin, W., Song, Y.: Integrating unmanned aerial vehicle-derived habitat metrics with movement persistence and species distribution models to characterize seasonal habitat use of invasive turtles in urban wetlands. Global Ecology and Conservation p. e04430 (2026)

29. Koch, F.H.: Considerations regarding species distribution models for forest insects. Agricultural and Forest Entomology 23(4), 393–399 (2021)

30. Koma, Z., Seijmonsbergen, A.C., Grootes, M.W., Nattino, F., Groot, J., Sierdsema, H., Foppen, R.P., Kissling, W.D.: Better together? assessing different remote sensing products for predicting habitat suitability of wetland birds. Diversity and distributions 28(4), 685–699 (2022)

31. Lauzeral, C., Grenouillet, G., Brosse, S.: Spatial range shape drives the grain size effects in species distribution models. Ecography 36(7), 778–787 (2013)

32. Levin, S.A.: The problem of pattern and scale in ecology: the robert h. macarthur award lecture. Ecology 73(6), 1943–1967 (1992)

33. Lipman, Y., Chen, R.T., Ben-Hamu, H., Nickel, M., Le, M.: Flow matching for generative modeling. arXiv preprint arXiv:2210.02747 (2022)

34. Liu, X., Frey, J., Munteanu, C., Denter, M., Koch, B.: Tree species diversity mapping from spaceborne optical images: The effects of spectral and spatial resolution. Remote Sensing in Ecology and Conservation 10(4), 463–479 (2024)

35. Lorieul, T., Cole, E., Deneu, B., Servajean, M., Bonnet, P., Joly, A.: Overview of geolifeclef 2022: Predicting species presence from multi-modal remote sensing, bioclimatic and pedologic data. In: CLEF (Working Notes). CEUR-WS (2022)

36. Loshchilov, I., Hutter, F.: Decoupled weight decay regularization. arXiv preprint arXiv:1711.05101 (2017)

37. Manzoor, S.A., Griffiths, G., Lukac, M.: Species distribution model transferability and model grain size–finer may not always be better. Scientific reports 8(1), 7168 (2018)

38. Mao, A., Mohri, M., Zhong, Y.: Cross-entropy loss functions: Theoretical analysis and applications. In: International conference on Machine learning. pp. 23803–23828. pmlr (2023)

39. Moudrỳ, V., Keil, P., Gábor, L., Lecours, V., Zarzo-Arias, A., Barták, V., Malavasi, M., Rocchini, D., Torresani, M., Gdulová, K., et al.: Scale mismatches between predictor and response variables in species distribution modelling: A review of practices for appropriate grain selection. Progress in Physical Geography: Earth and Environment 47(3), 467–482 (2023)

40. Müllerová, J., Brůna, J., Bartaloš, T., Dvořák, P., Vítková, M., Pyšek, P.: Timing is important: Unmanned aircraft vs. satellite imagery in plant invasion monitoring. Frontiers in Plant Science 8, 887 (2017)

41. Nagendra, H., Rocchini, D., Ghate, R., Sharma, B., Pareeth, S.: Assessing plant diversity in a dry tropical forest: Comparing the utility of landsat and ikonos satellite images. Remote sensing 2(2), 478–496 (2010)

42. Okabayashi, A., Audebert, N., Donike, S., Pelletier, C.: Cross-sensor super-resolution of irregularly sampled sentinel-2 time series. In: 2024 IEEE/CVF Conference on Computer Vision and Pattern Recognition Workshops (CVPRW). pp. 502–511. IEEE (2024)

43. O’Neill, R.V.: A hierarchical concept of ecosystems. No. 23 in Monographs in Population Biology, Princeton University Press (1986)

44. Peterman, W.E., Connette, G.M., Semlitsch, R.D., Eggert, L.S.: Ecological resistance surfaces predict fine-scale genetic differentiation in a terrestrial woodland salamander. Molecular ecology 23(10), 2402–2413 (2014)

45. Picek, L., Botella, C., Servajean, M., Leblanc, C., Palard, R., Larcher, T., Deneu, B., Marcos, D., Estopinan, J., Bonnet, P., et al.: Overview of geolifeclef 2024: Species composition prediction with high spatial resolution at continental scale using remote sensing. In: CLEF 2024-Working Notes of the 25th Conference and Labs of the Evaluation Forum). CEUR-WS (2024)

46. Raitif, J., Plantegenest, M., Roussel, J.M.: From stream to land: Ecosystem services provided by stream insects to agriculture. Agriculture, ecosystems & environment 270, 32–40 (2019)

47. Raitif, J., Roussel, J.M., Olmos, M., Piscart, C., Plantegenest, M.: Assessing spatial deposition of aquatic subsidies by insects emerging from agricultural streams. Science of the Total Environment 837, 155686 (2022)

48. Ronneberger, O., Fischer, P., Brox, T.: U-net: Convolutional networks for biomedical image segmentation. In: International Conference on Medical image computing and computer-assisted intervention. pp. 234–241. Springer (2015)

49. Ruwaimana, M., Satyanarayana, B., Otero, V., M. Muslim, A., Syafiq A, M., Ibrahim, S., Raymaekers, D., Koedam, N., Dahdouh-Guebas, F.: The advantages of using drones over space-borne imagery in the mapping of mangrove forests. PloS one 13(7), e0200288 (2018)

50. Selvaraju, R.R., Cogswell, M., Das, A., Vedantam, R., Parikh, D., Batra, D.: Grad-cam: Visual explanations from deep networks via gradient-based localization. In: Proceedings of the IEEE international conference on computer vision. pp. 618–626 (2017)

51. Seoane, J., Bustamante, J., Díaz-Delgado, R.: Are existing vegetation maps adequate to predict bird distributions? Ecological Modelling 175(2), 137–149 (2004), https://www.sciencedirect.com/science/article/pii/S0304380003004733

52. Silva, D.P., Spigoloni, Z.A., Camargos, L.M., de Andrade, A.F.A., De Marco Jr, P., Engel, M.S.: Distributional modeling of mantophasmatodea (insecta: Notoptera): a preliminary application and the need for future sampling: Dp silva et al. Organisms Diversity & Evolution 16(1), 259–268 (2016)

53. Šímová, P., Moudrỳ, V., Komárek, J., Hrach, K., Fortin, M.J.: Fine scale water-body data improve prediction of waterbird occurrence despite coarse species data. Ecography 42(3), 511–520 (2019)

54. Teillet, C., Pottier, H., Devillers, R., Defossez, A., Catry, T., Kerr, A., Jean, F., L’ambert, G., LeDoeuff, N., Roux, E.: Characterizing urban landscapes using very-high resolution satellite imagery to predict ae. albopictus larval presence probability in public spaces. Plos one 20(11), e0335794 (2025)

55. Teng, M., Elmustafa, A., Akera, B., Bengio, Y., Radi, H., Larochelle, H., Rolnick, D.: Satbird: a dataset for bird species distribution modeling using remote sensing and citizen science data. In: Oh, A., Naumann, T., Globerson, A., Saenko, K., Hardt, M., Levine, S. (eds.) Advances in Neural Information Processing Systems. vol. 36, pp. 75925–75950. Curran Associates, Inc. (2023). 10.52202/075280-3317, https://proceedings.neurips.cc/paper_files/paper/2023/file/ef7653bbc4655305efb89a32362e332a-Paper-Datasets_and_Benchmarks.pdf

56. Theron, K.J., Pryke, J.S., Latte, N., Samways, M.J.: Mapping an alien invasive shrub within conservation corridors using super-resolution satellite imagery. Journal of Environmental Management 321, 116023 (2022). 10.1016/j.jenvman.2022.116023, https://www.sciencedirect.com/science/article/pii/S0301479722015961

57. van Tiel, N., Zbinden, R., Vanalli, C., Kellenberger, B., Pellissier, L., Tuia, D.: Multi-scale neural networks enhance species distribution modelling across predictors and taxonomic groups. Ecological Informatics 96, 103799 (2026). 10.1016/j.ecoinf.2026.103799, https://www.sciencedirect.com/science/article/pii/S1574954126002050

58. Vogt, P., Riitters, K.H., Estreguil, C., Kozak, J., Wade, T.G., Wickham, J.D.: Mapping spatial patterns with morphological image processing. Landscape Ecology 22(2), 171–177 (2007). https://doi.org/10.1007/s10980-006-9013-2, 10.1007/s10980-006-9013-2

59. Wen, K., Luo, Y., Rao, Z., Wu, A., Song, Q., Zhao, Y., Lyu, Y., Zou, Q., Chen, M., Niu, K., et al.: Use of drone-derived high-resolution elevation data to improve species distribution models for a primate in mountainous areas. Conservation Biology p. e70274 (2026)

60. Wiens, J.A.: Spatial scaling in ecology. Functional ecology 3(4), 385–397 (1989)

61. Zhang, H., Cisse, M., Dauphin, Y.N., Lopez-Paz, D.: mixup: Beyond empirical risk minimization. arXiv preprint arXiv:1710.09412 (2017)

